## Supplementary data for "Reassessing the substrate specificities of the major *Staphylococcus aureus* peptidoglycan hydrolases lysostaphin and LytM"

Supplementary material

Lina Antenucci[1], Salla Virtanen[2], Chandan Thapa[1], Minne Jartti[1,#], Ilona Pitkänen[1], Helena Tossavainen[1], Perttu Permi[1,2,3,*]

[1]*Department of Biological and Environmental Science, Nanoscience Center, University of Jyvaskyla, P.O. Box 35, FI-40014 Jyvaskyla, Finland*

[2]*Institute of Biotechnology, Helsinki Institute of Life Science, University of Helsinki, Helsinki, Finland*

[3]*Department of Chemistry, Nanoscience Center, University of Jyvaskyla, P.O. Box 35, FI-40014 Jyvaskyla, Finland*

[#]*Present address: Faculty of Medicine and Health Technology, Tampere University, Tampere, Finland*

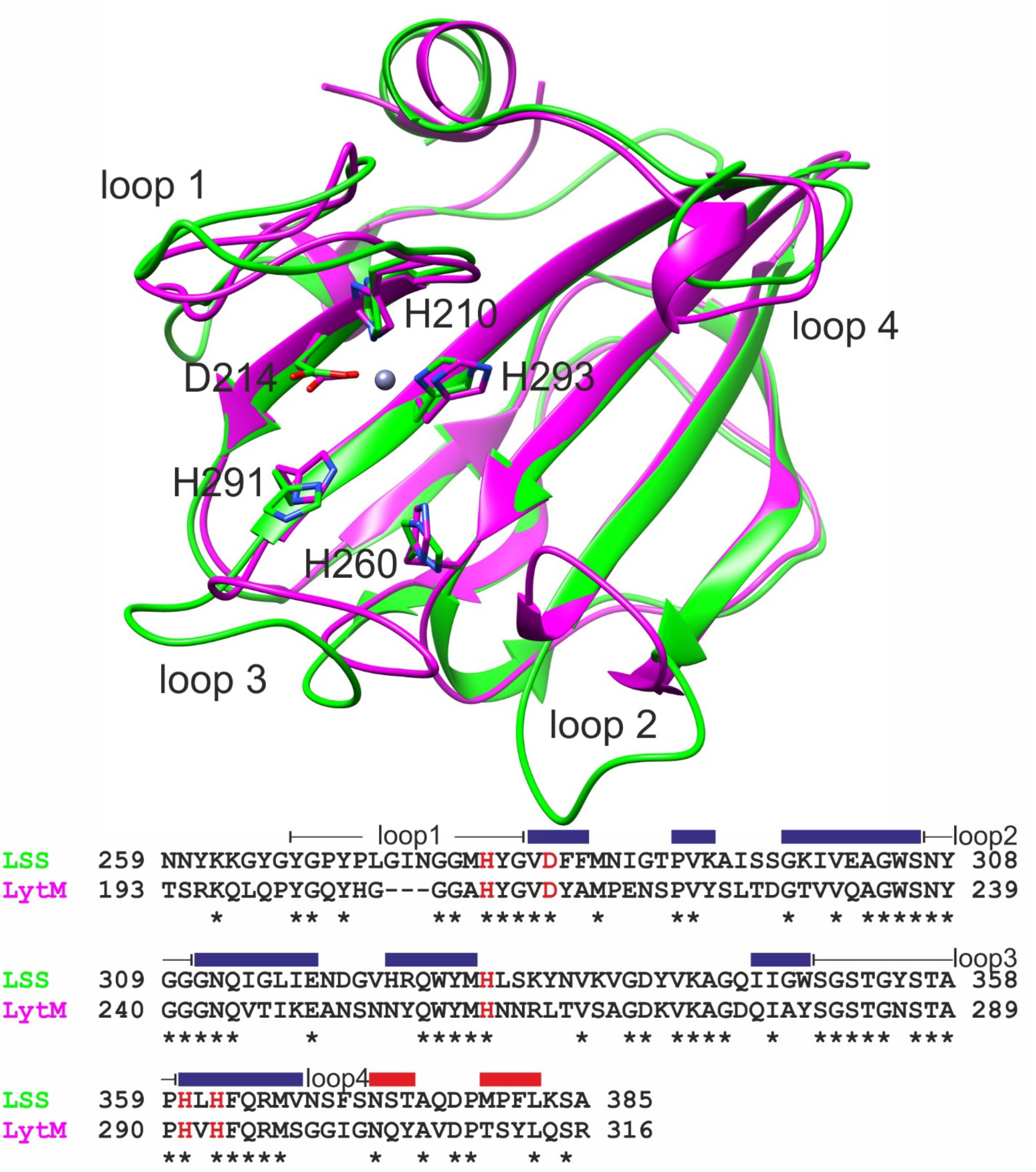


**Fig. S1**. Overlaid structures of the catalytic domains of LSS (green, PDB ID 5NMY) and LytM (magenta, PDB ID 2B13) as well as their aligned amino acid sequences. Side chains of zinc-coordinating and catalytic residues are shown in sticks and highlighted in red in the sequence alignment. Loops surrounding the catalytic groove are numbered from 1 to 4, and together with secondary structures (β strands in blue and α helices in red) shown for LytM above the sequence.

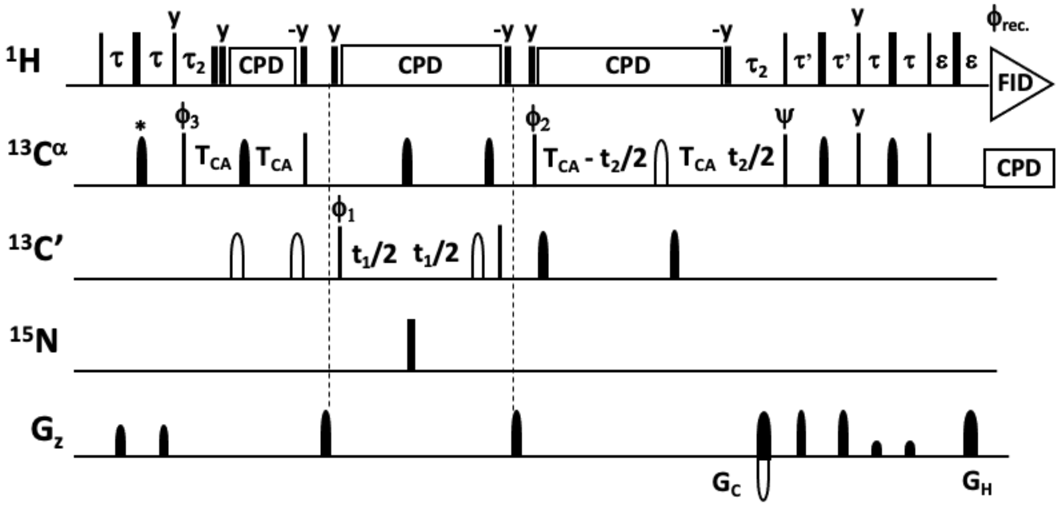


**Fig. S2.** Glycine-optimized HCACO experiment for correlating $^{1}$Hα, $^{13}$Cα, $^{13}$CO resonances. Narrow and wide filled bars on $^{1}$H and $^{15}$N channels correspond to rectangular 90° and 180° pulses, respectively, applied with phase x unless otherwise stated. All $^{13}$C pulses are band-selective shaped pulses, denoted by filled narrow bars (90°) and filled and unfilled half ellipsoids (180°). Pulses denoted with unfilled bars are applied on-resonance. The $^{1}$H, $^{15}$N, $^{13}$CO, and $^{13}$Cα carrier positions are 4.7 (water), 118 (center of $^{15}$N spectral region), 174 ppm (center of $^{13}$CO spectral region), and 54 ppm (center of $^{13}$Cα spectral region). The $^{13}$C carrier is initially set to the middle of $^{13}$Cα region (45 ppm) and shifted to $^{13}$CO region (174 ppm) prior to 90° $^{13}$CO pulse $\phi_1$. All band-selective 90° and 180° pulses for $^{13}$Cα (45 ppm) and $^{13}$CO (174 ppm) have the shape of Q5 and Q3 [1] and duration of 240.0 μs and 192.0 μs at 800 MHz, respectively. The Waltz-65 sequence [2] with strength of 4.17 kHz was employed to decouple $^{1}$H spins. The GARP [3,4] with field strength of 4.55 kHz was used to decouple $^{13}$C during acquisition. Delay durations: $\tau = 1/(4J_{HC}) \sim 1.7$ ms; $\tau' = 0.85$ ms; $\tau_2 = 1.7$ ms (optimized for glycine residues) or $\tau_2 = 2.2 – 2.6$ ms (for observing both glycine and non-glycine residues); $\varepsilon$ = duration of $G_H$ + field recovery ~1.2 ms; $2T_{CA} = 1/(2J_{C\alpha CO}) \sim 9.5$ ms. Maximum $t_2$ is restrained $t_{2,max} < 2.0*(T_{CA} - \tau_2)$. Frequency discrimination in $^{13}$CO dimension is obtained using the States-TPPI protocol [5] applied to $\phi_1$, whereas the quadrature detection in $^{13}$Cα dimension is obtained using the sensitivity-enhanced gradient selection [6,7]. The echo and anti-echo signals in $^{13}$Cα dimension are collected separately by inverting the sign of the $G_C$ gradient pulse together with the inversion of ψ, respectively. Phase cycling: $\phi_1$ = x, -x; $\phi_2$ = 2(x), 2(-x); $\phi_3$ = 4(x), 4(-x); ψ = x; rec. = x, 2(-x), x, -x, 2(x), -x. Gradient strengths (% of max G/cm) and durations (ms) for the coherence selection: $G_C$ = 80.0 %, 1.0 ms; $G_H$ = 20.1 % 1.0 ms. The pulse sequence code and parameter file for Bruker Avance system are available from authors upon request.

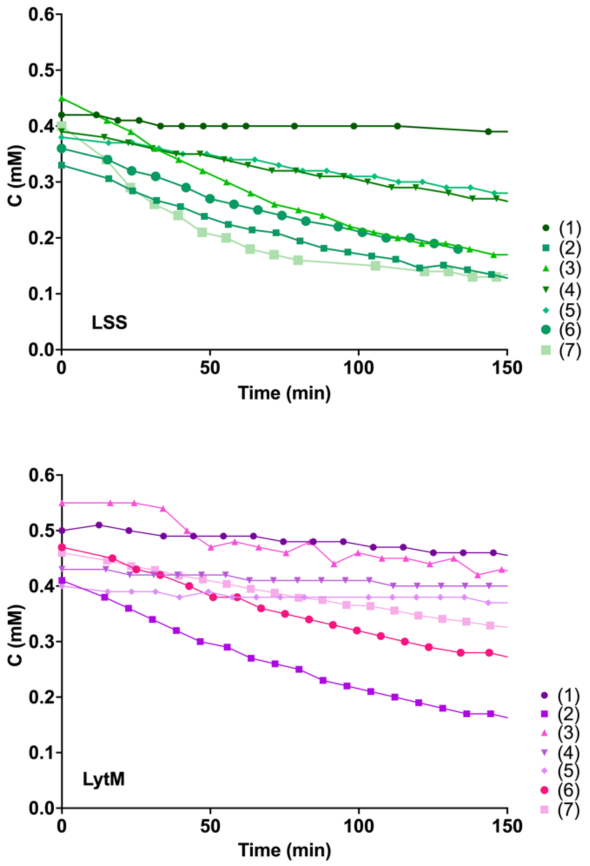


**Fig. S3.** Hydrolysis of peptides **1**-**7** by LSS (green) and LytM (magenta). Plots of decaying substrate concentration in function of reaction time.

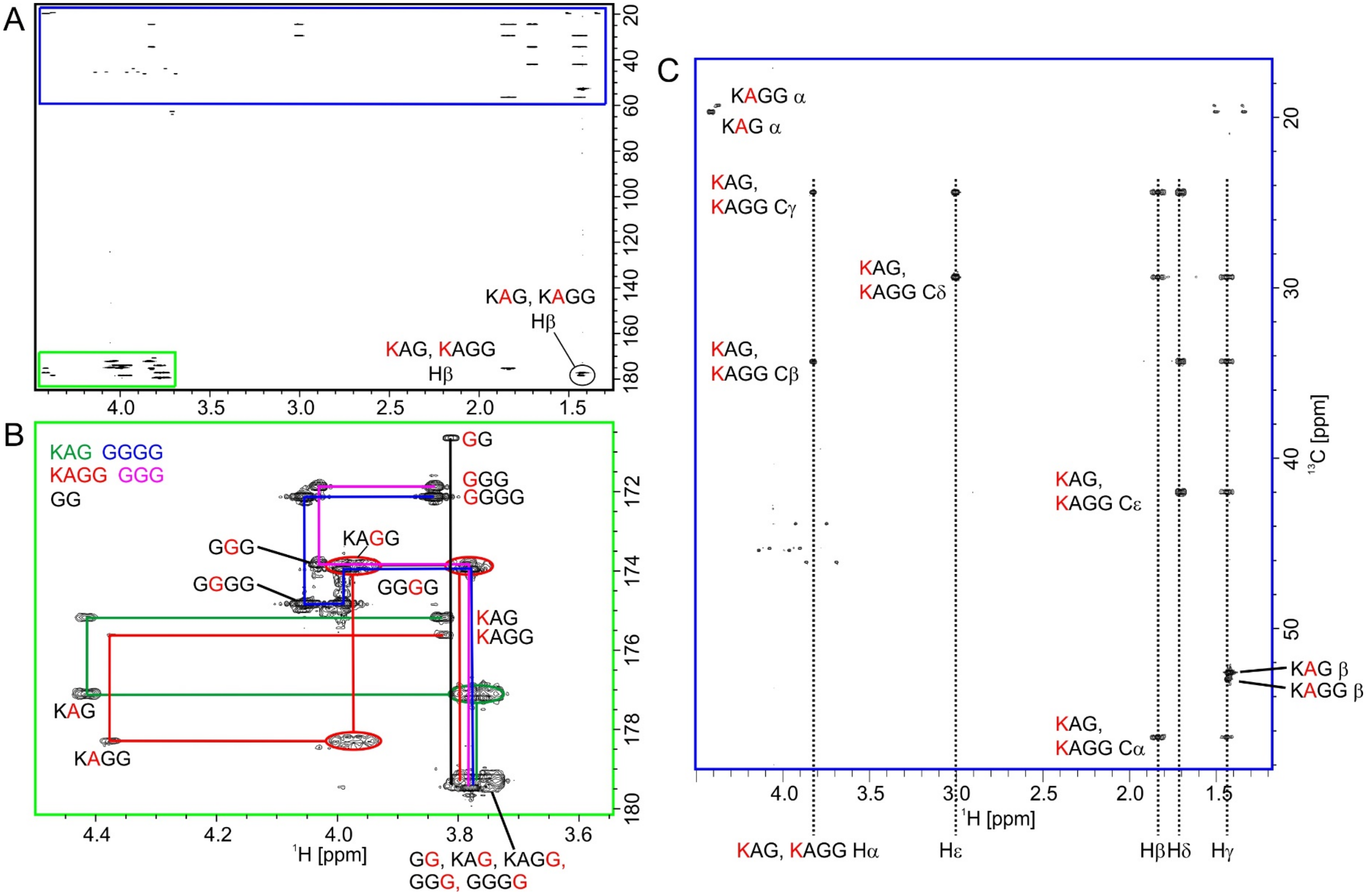


**Fig. S4.** Identification of products in the hydrolysis reaction of peptide **2** by LSS with a $^{13}$C-HMBC spectrum. In **A)** is shown the full spectrum. Expansions from the boxed regions are shown in B) corresponding to the Hα–carbonyl carbon region and **C)** the aliphatic region. In the former, inter-residual connectivities of the five products are shown with lines of different colors. Diglycine is a secondary product from hydrolysis of tetraglycine.

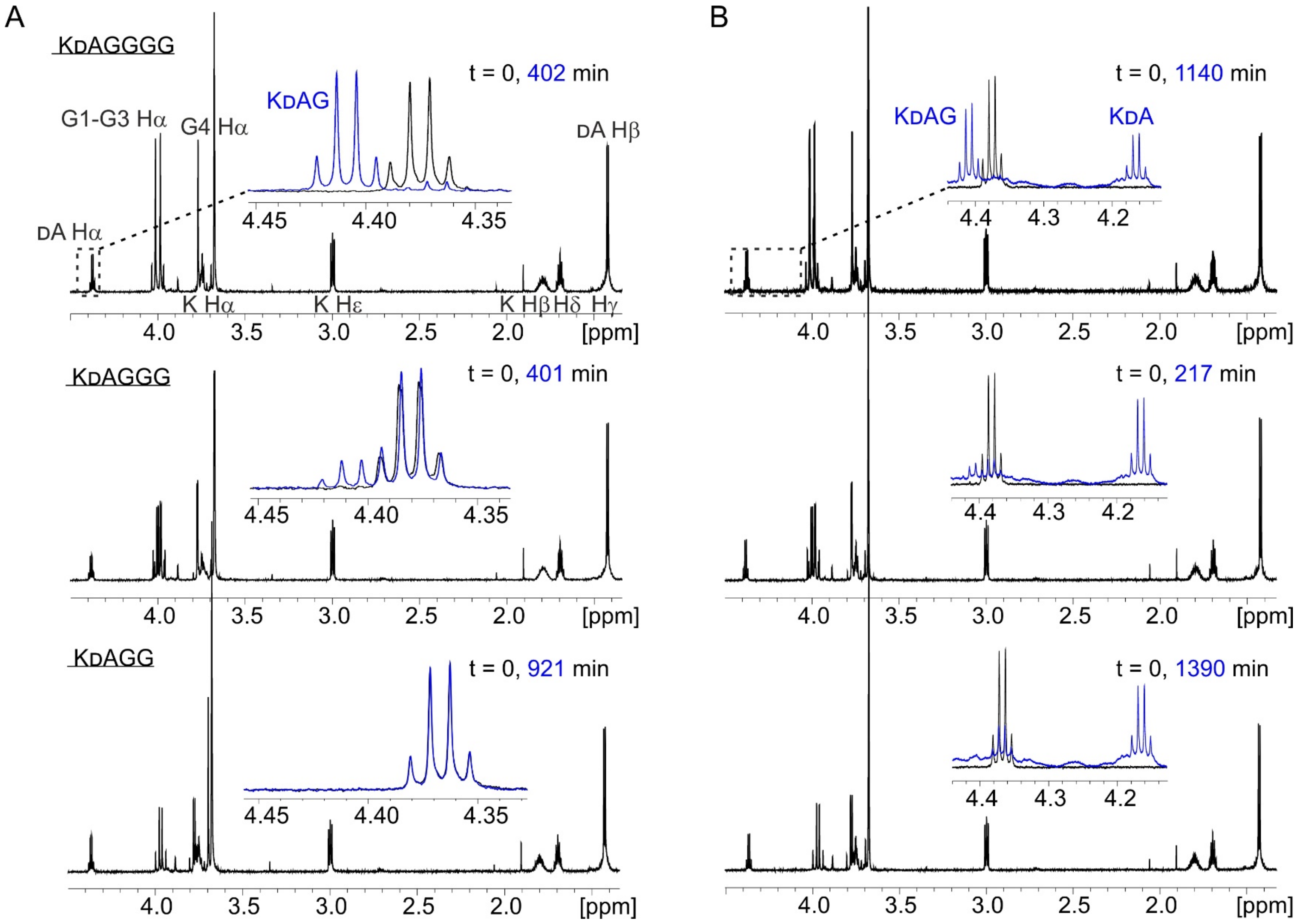


**Fig. S5.** Hydrolysis of peptides KDAGGGG (**12**), KDAGGG (**13**) and KDAGG (**14**) by LSS (**A**) and LytM (**B**). In the insets are shown overlays of the boxed DA Hα region of the $^{1}$H spectra at two time points of the hydrolysis reactions.

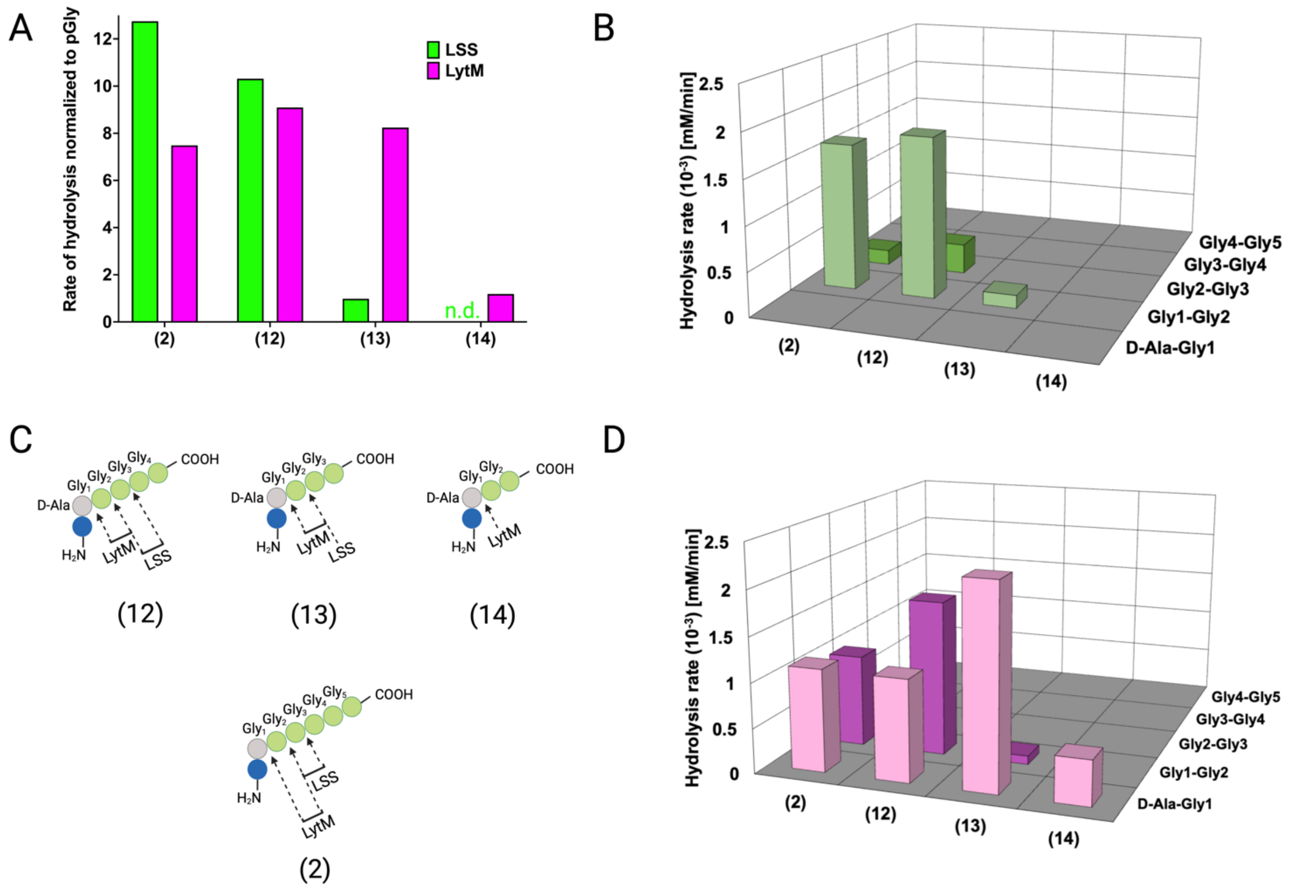


**Fig. S6.** Hydrolysis of PG fragments **12-14** by LSS and LytM. **A)** Rate of substrate hydrolysis with respect to pGly. Shown are comparative rates for the substrate **2** and **12-14**, having five, four, three and glycines cross-linked to D-Ala, respectively. **B)** Rates of scissile bond hydrolysis in substrates **2**, and **12-14** for LSS. **C)** Bonds cleaved by LytM and LSS in the PG fragments **2**, **12-14**. **D)** Rates of scissile bond hydrolysis in substrates **2**, and **12-14** for LytM.

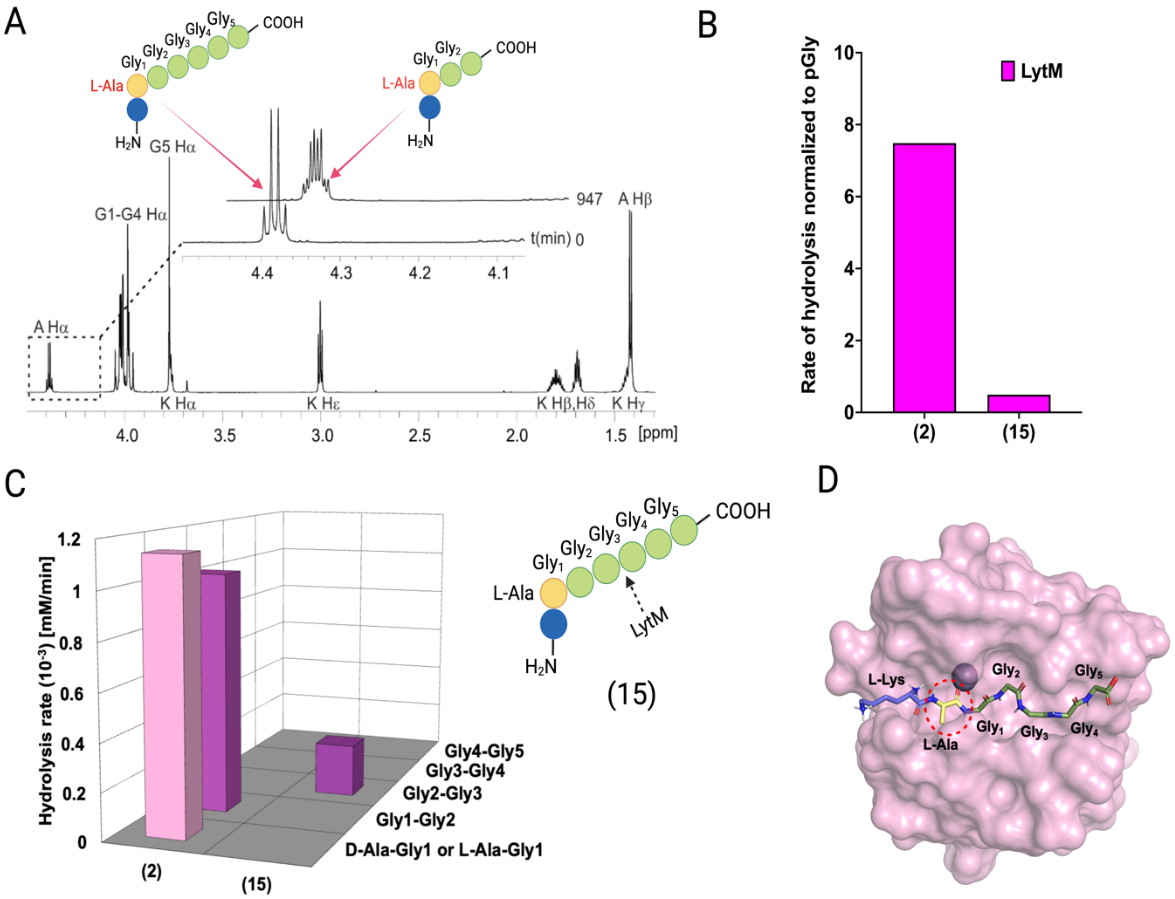


**Fig. S7.** Hydrolysis of peptide KLAGGGGG (**15**) by LytM. **A)** Contrary to cleavages between Ala-Gly$_1$ and Gly$_1$-Gly$_2$ in KDAGGGGG, LytM now uniquely cleaves between Gly$_2$ and Gly$_3$. The Hα peak at time point 947 minutes is composed of remaining substrate Ala Hα and Ala Hα from KLAGG. **B)** The overall rate of substrate hydrolysis of KLAGGGGG mimics that of pGly and is much slower than hydrolysis of the PG fragment **2**. **C)** Scissile bond (Gly2-Gly3) is the same as for pGly (**1**). **D)** L-Ala-Gly$_1$ bond hydrolysis is hindered by steric clash of Ala methyl group and loop 3.

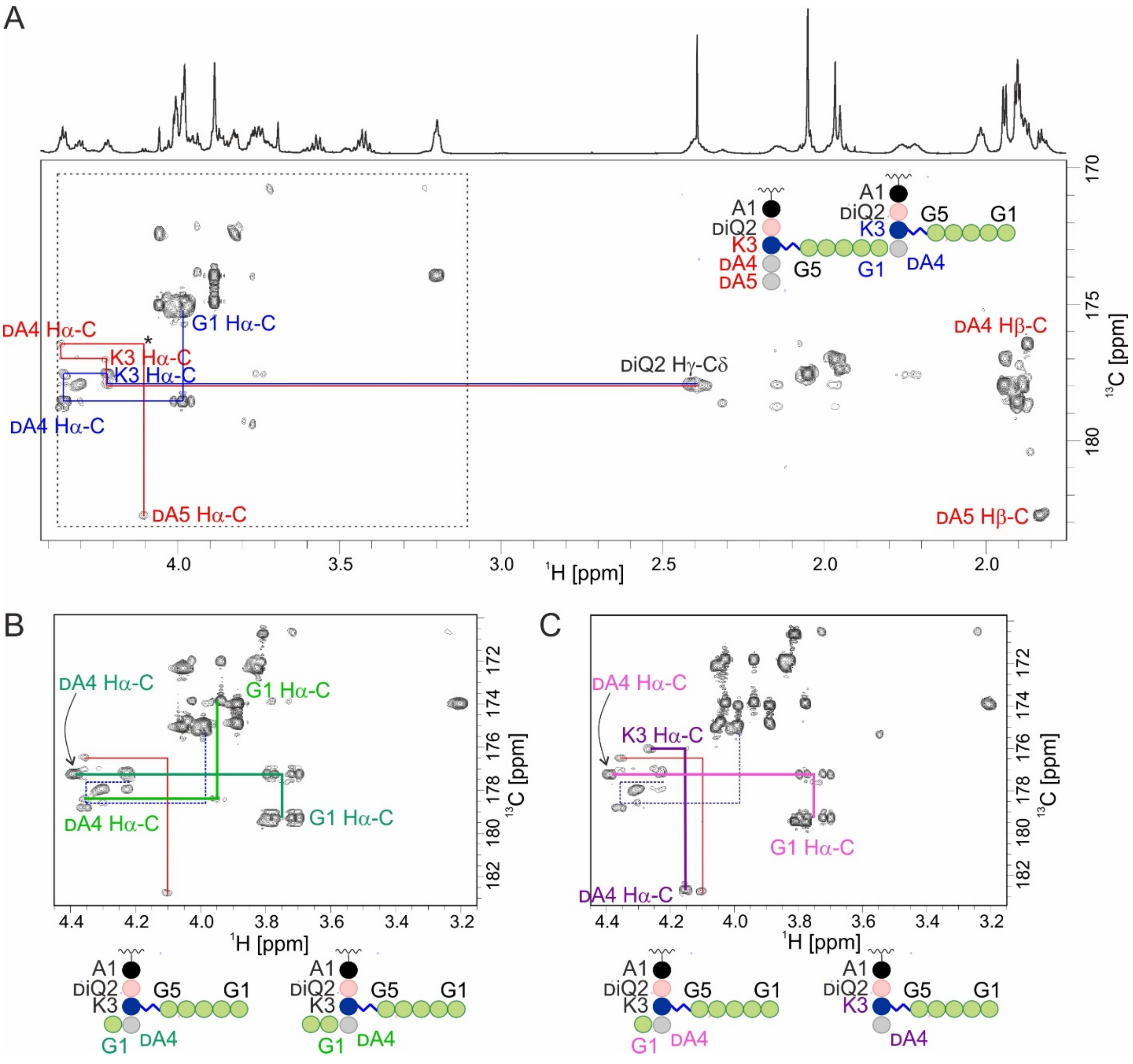


**Fig. S8.** Hydrolysis of muropeptides extracted from *S. aureus* USA300 sacculus by LSS and LytM. **A)** The carbonyl carbon region of a $^{13}$C-HMBC spectrum acquired from muropeptides prior to enzyme addition. Inter-residual connectivities and identities are shown for some of the peaks. Color codes match between the latter and the schematic structure of a PG monomer in the upper right corner. The asterisk marks a peak seen at lower contour levels. **B)** The boxed region after reaction with LSS. LSS cleaves the bond between glycines 1 and 2 and a characteristic quartet-like peak pattern of an Ala-linked C-terminal glycine Hα appears (dark green). Based on signal intensities here and those in the $^{13}$C-HMBC of reaction products of hydrolysis of **2** by LSS (Fig. S3), cleavage between glycines 2 and 3 occurs more seldom (bright green). **C)** The boxed region after reaction with LytM. Similar to the LSS-reaction, the G1-G2 bond cleavage product is present (light purple). Additionally, the peaks of a Lys-linked C-terminal alanine appear (dark purple).

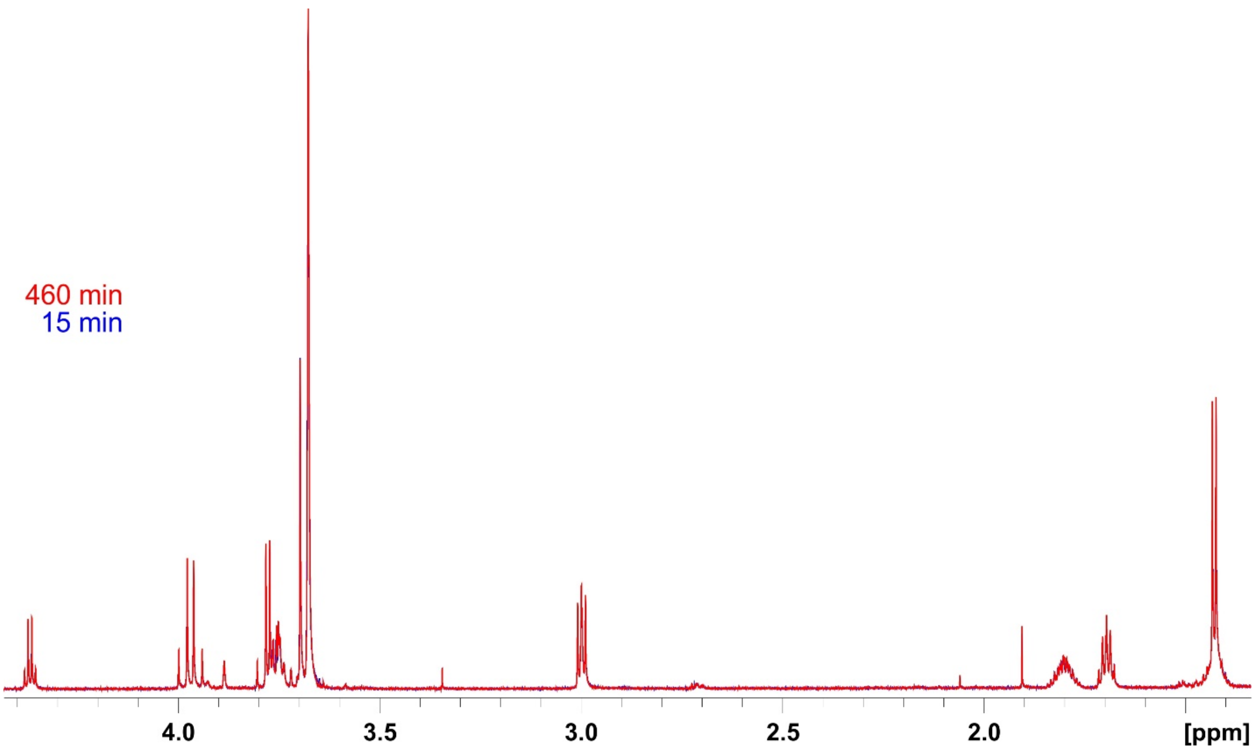


**Fig. S9**. Assessment of the stability of the 800 MHz $^{1}$H NMR spectrometer during the relatively long periods of acquisition of kinetics spectra. Shown are overlaid $^{1}$H spectra at the start and end points of the hydrolysis reaction of peptide **14** by LSS. This PG fragment is not hydrolysed by LSS. Substrate concentration was 0.4 mM and that of LSS 2 μM.

### Table S1. Synthetic PG fragments used in this study

| | Description | PG fragment |
|---|---|---|
| 1 | pentaGly (pGly) | $H_2N$-GGGGG-COOH |
| 2 | pGly N-term linked to D-Ala-Lys | $H_2N$-KDA-GGGGG-COOH |
| 3 | pGly N-term linked to tetrapeptide stem | $H_2N$-ADiQKDA-GGGGG-COOH |
| 4 | pGly C-term linked to Nζ of Lys tetrapeptide stem | $H_2N$-ADiQK(Nζ-GGGGG-$NH_2$)DA-COOH |
| 5 | pGly C-term linked to Nζ of Lys in pentapeptide stem | $H_2N$-ADiQK(Nζ-GGGGG-$NH_2$)DADA-COOH |
| 6 | pGly C-term linked to Nζ of Lys in tetrapeptide stem and N-term to D-Ala-Lys | $H_2N$-ADiQK(Nζ-GGGGG-DAK-$NH_2$)DA-COOH |
| 7 | pGly C-term linked to Nζ of Lys in pentapeptide stem and N-term to tetrapeptide stem | $H_2N$-ADiQK(Nζ-GGGGG-DAKDiQA-$NH_2$)DADA-COOH |
| 8 | triGly C-term linked to Nζ of Lys in pentapeptide stem and N-term to tetrapeptide stem | $H_2N$-ADiQK(Nζ-GGG-DAKDiQA-$NH_2$)DADA-COOH |
| 9 | monoGly C-term linked to Nζ of Lys in pentapeptide stem and N-term to tetrapeptide stem | $H_2N$-ADiQK(Nζ-G-DAKDiQA-$NH_2$)DADA-COOH |
| 10 | GSGGG N-term linked to D-Ala-L-Lys | $H_2N$-KDA-GSGGG-COOH |
| 11 | GGSGG N-term linked to D-Ala-L-Lys | $H_2N$-KDA-GGSGG-COOH |
| 12 | tetraGly N-term linked to D-Ala-L-Lys | $H_2N$-KDA-GGGG-COOH |
| 13 | triGly N-term linked to D-Ala-L-Lys | $H_2N$-KDA-GGG-COOH |
| 14 | diGly N-term linked to D-Ala-L-Lys | $H_2N$-KDA-GG-COOH |
| 15 | pGly N-term linked to L-Ala-L-Lys | $H_2N$-KA-GGGGG-COOH |
